# Performance Evaluation of Markerless 3D Skeleton Pose Estimates with Pop Dance Motion Sequence

**DOI:** 10.1101/2020.04.15.010702

**Authors:** Rollyn T. Labuguen, Wally Enrico M. Ingco, Salvador Blanco Negrete, Tonan Kogami, Tomohiro Shibata

**Affiliations:** Graduate School of Life Science and Systems Engineering, Kyushu Institute of Technology, Kitakyushu City, Fukuoka, Japan; Electronics, Computer and Communications Engineering, Ateneo de Manila University, Quezon City, Philippines

**Keywords:** markerless human pose estimation, OpenPose evaluation, motion capture, pop dance

## Abstract

The evaluation of markerless pose estimation performed by OpenPose has been getting much attention from researchers of human movement studies. This work aims to evaluate and compare the output joint positions estimated by the OpenPose with a marker-based motion-capture data recorded on a pop dance motion. Although the marker-based motion capture can accurately measure and record the human joint positions, this particular set-up is expensive. The framework to compare the outputs of the markerless method to the ground truth marker-based joint remains unknown, especially for complex body motion. Synchronization, camera calibration, and 3D reconstruction by fusing the outputs of the markerless method (OpenPose) are discussed. In this case study, the comparison results illustrate that even if the markerless method expects to fail when the subject’s body parts are self-occluded, the average magnitude errors for each key points are less than 700 mm.

## I. Introduction

The genre for pop dance (also known as robotic dancing) is rapidly growing and has been introduced and popularized in different cultures around the world. This is a dance style where the dancer moves and imitates a mannequin with a mechanical stutter along with pop music. It usually involves complex and agile steps wherein dancers are training and practicing these steps frequently to gain a smooth transition of the sequences. Although dancing is mostly deemed as an expression of art, human behavior scientists, biomechanics researchers, and gaming application developers have interests in quantifying and digitizing the dancers’ movements.

Motion capture (MoCap) systems have been used to record and track the movements of human joint positions. Though these motion capture systems’ accuracy rely on correct markers placement and calibrated setup, these methods can be intrusive to the natural behavior of the subject’s dancing. MoCap is also expensive, and sometimes markers tend to be detached from the body joint when the subject performs complex motions. As such, the markerless methods are introduced by using deep learning techniques and huge human datasets. OpenPose [1], one of the most popular markerless human pose estimation method, is easy-to-use and is applicable to both videos and images. OpenPose is capable of pose estimation for multiple subjects, however, its accuracy and robustness for complex motions are yet to be tested. Comparison between OpenPose and MoCap system on the 3D space has only been performed on a dataset that includes simple actions such as walking, jumping and throwing [2]. Hence, we want to more quantitatively assess whether the OpenPose would be able to handle fast and complex dance movements, as this markerless method will not require a tedious setup and is not intrusive of the subject’s motion.

## II. Related Works

### A. Marker-based Motion Capture Studies for Dancing

Motion capture studies for dancing have been done and collected to enable digitization and preservation of the performances. In the book [4] “Dance Notations and Robot Motion”, numerous publications were published regarding the use of motion capture systems including the study to analyze Tango dance which is a slow dance style sequence [3]. A similar study using motion capture data and Laban Movement Analysis LMA [4], was applied to folk dance by Aristidou et al. [5]. The folk dancing has beats and the researchers have implemented a virtual simulator for demonstrating the dance style wherein the users can preview the dance steps performed by a 3D avatar.

### B. Markerless Skeleton Pose Estimates Evaluation

Digitizing the whole-body motions of human dancer/s, the motion capture system shows its high reliability in terms of tracking a set of external markers. But these markers often affect the dancer’s dynamic motion and its performance during the dance motion capturing sessions. Yejin Kim [6] introduced a markerless motion capture and composition system for a ballet dance motion that utilizes multiple RGB and depth sensors. A similar motivation is seen in [7] but this study concentrates on K-Pop dance and the researchers develop a 2D markerless pose estimation that is invariant to full-body rotation and self-occlusion. Their framework uses ridge data and data pruning to estimate the human pose and utilizes expert dance database to evaluate a novice or dance learners’ steps.

Our work relates and differs from the previous studies by the following contributions:

- Captured expert pop dance data recorded simultaneously with multiple RGBD sensors and application of the existing markerless approach to estimate 3D dance pose.
- Performance evaluation of OpenPose estimation in three-dimensional space with the marker-based system data.

## III. Experimental Setup

For the recording of the dataset, a professional popping dancer participated in the motion-capture study. The physical characteristics of the participant were as follows: 174 cm height, 68kg of body mass and 36 years of age.

The dancer performed freestyle popping with no choreography. Although six dancing sessions were recorded, for this study we will only focus on one recorded session. The selected session has a duration of one minute, which includes a variety of complex and agile dance steps. The session was recorded using five devices, a handicam, three Intel RealSense D435, and the MAC-3D. Three different sensors recorded the dance session simultaneously. To synchronize and calibrate the multiple recordings, the dancer performed a T-pose as illustrated in Figure 1. Then the videos by the Intel RealSenes and handicam were analyzed by OpenPose to extract 2D markerless keypoint position estimates. Using processing as explained in Section IV, 3D pose estimates were calculated.

**Fig. 1.**
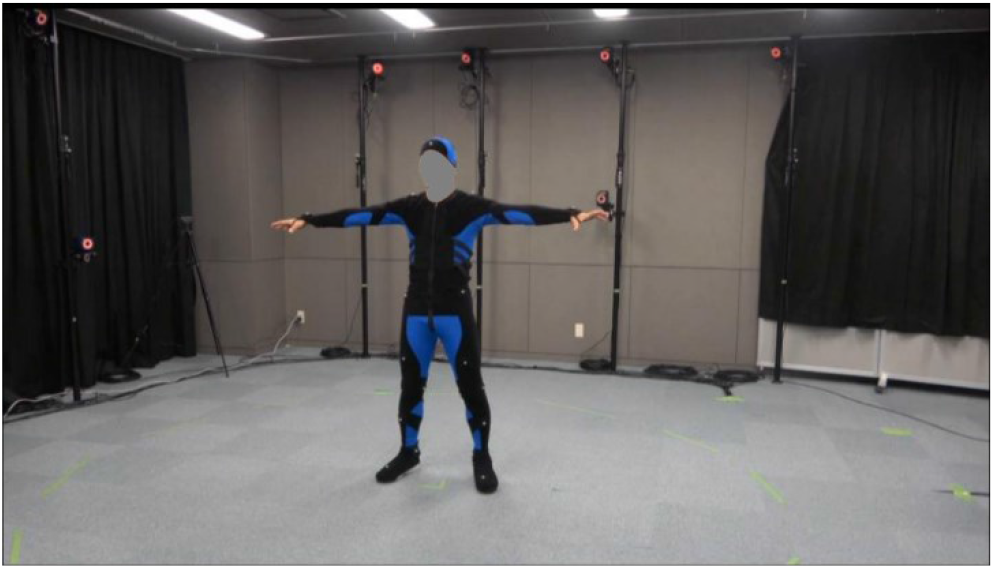
Example of a T-pose of the dancer

### A. Sensors for Motion Acquisition

#### 1) Handicam and Realsense Cameras

To record in 2D the handicap SONY FDR-AX45 was used. Three Intel Realsense D435 were also used to capture the dance moves in RGBD (.rosbag) format.

**Table I.** Recording Parameters

|  | Handicam | Intel Realsense D435 |
| --- | --- | --- |
| Video Resolution | 720p | 720p |
| Frame Rate | 30 fps | 30 fps |

OpenPose v1.5.1 was used to analyze the videos using the default settings utilizing the COCO model, with *xy* coordinates of twenty-five (25) body features as outputs.

#### 2) Motion Capture Marker-based Setup

Twenty-nine reflective markers are attached to the subject’s body as landmarks to be tracked by an 18-camera Kestrel 1300 Nac Image Technology Inc., at a sampling rate of 100 Hz. The marker placements followed the modified Helen Hayes marker illustrated by [8]. The accompanying software used is Cortex for preprocessing, capturing, and post-processing of datasets. Post-processing of an experimenter takes almost an hour to correctly annotate and connect joint markers properly. We also utilized OpenSim for quick visualization.

### B. Multi-camera Calibration

The motion capture system was calibrated initially using the conventional wand method for dynamic calibration and the L-frame for static. Since there is also an independent multiple RGB-D camera setup placed during the dance experiment, the calibration for this setup must be made. For the depth sensors and handicam calibration, we base the calibration method using the markers when the dancer made a T-pose. By using the pin-hole method equation for each camera, we could compute for **P** using (1).

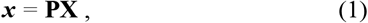

where ***x*** contains the 2D image point coordinate of a marker, **X** is the corresponding 3D world point from the MoCap data, **P** is the camera matrix containing the camera intrinsic and extrinsic parameters. Generally, matrix **P** = K [R t] where K is a 3×3 matrix containing the focal length and principal point of the camera, R is a 3×3 rotation matrix and lastly, t represents 3×1 translation vector. [R t] encodes the orientation and position of the cameras with respect to a reference coordinate system. The reference coordinate system used is the global coordinate system based on the MoCap.

Figure 2 illustrates the derived and calibrated positions of the cameras from the T-pose world coordinate points. Black points are the body markers, triangle shapes refer to the cameras (Blue: handicam, Magenta: Intel Realsense Camera 1 looking at the back of the subject, Green: Intel Realsense Camera 2 looking at the side of the subject, Red: Intel Realsense Camera 3 looking at the front of the subject).

**Fig. 2.**
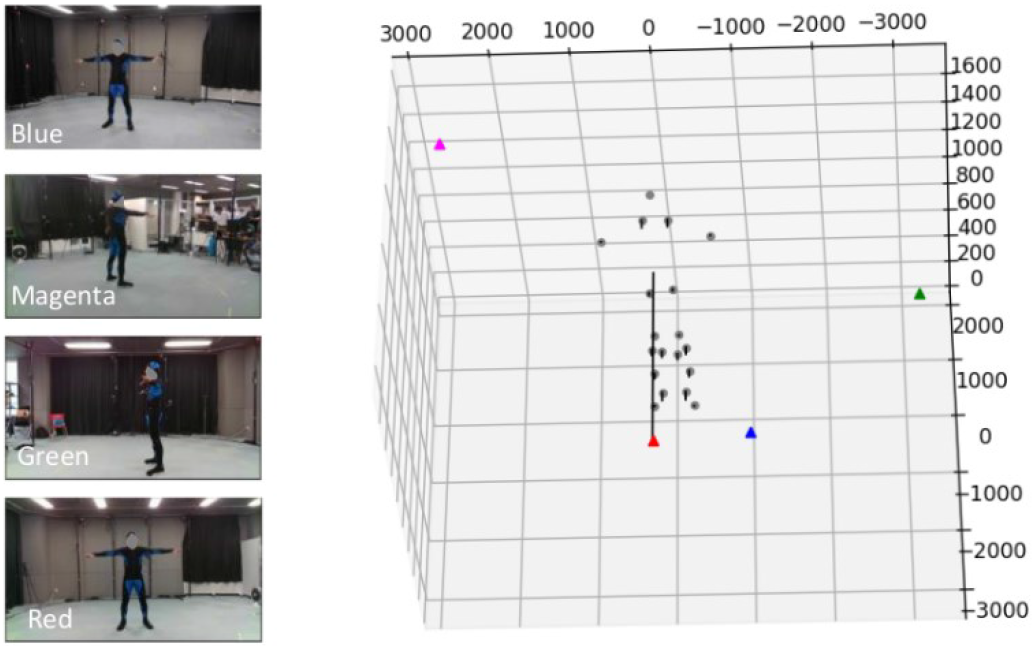
Handicam Position (Blue) and Intel Realsense Camera Positions

### C. Data Synchronization

The Handicam Camera, Realsense Camera, and MAC3D were all synchronized using the global time frame. The Realsense cameras’ frame rate is similar to the Handicam’s frame rate, valued at 30 fps. However, this frame rate does not equate with the frame rate of the MAC3D system. Each capture system produces a different number of frames at a certain time period making synchronizing a challenge. The MAC3D system will need to drop some frames in order to match the number of frames produced by the other two capture devices. Unfortunately, despite the frame-drops and matching of the frame numbers, delay of frames was still observed. The output video length from each camera is also different.

We synchronized the videos by using an editing software named OpenShot. OpenShot is a free Windows video editing software, with the timelining capability and a sensitivity of 0.01 seconds. Looking at a property that is common among the recording devices, time is only the invariant factor without issues such as frame drop. Thus, we selected a reference point in time to do synchronization, for example, when the subject performed the T-Pose. This T-Pose action can be observed in all video recordings. The video editor software OpenShot was used to align the videos as well as cut and trim accordingly.

For the MAC3D data, it was aligned by visualizing the pose of the subject using Opensim. The common point was visualized and adjustments on the .trc file were performed to match it with the other videos.

### D. Linear Regression for Predicting Additional Key Points to OpenPose

Some markers’ positions were not seen on the key points detection of OpenPose. We hypothesized that the positions of these markers not extracted by the OpenPose can be estimated using linear regression. For example, as to the Front Head, Rear Head, and the Top Head markers, their positions can assumingly have linear relationship with the OpenPose Nose key point. In the following figures, the data for the x-coordinates are shown.

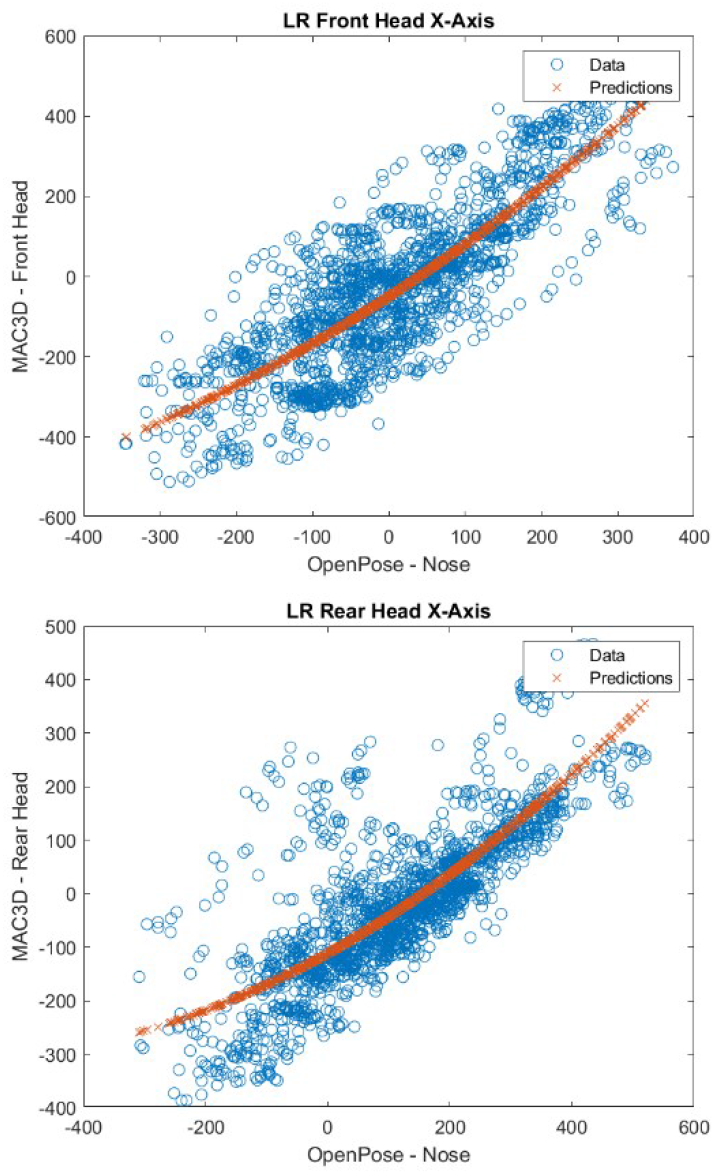

The x-axis of the OpenPose (Nose) was related to the x-axis of the MAC3D (Front, Top, Rear Head). Similar method was employed in the y and z axis. From the two variables, a regression model is created and fitted with the existing data.

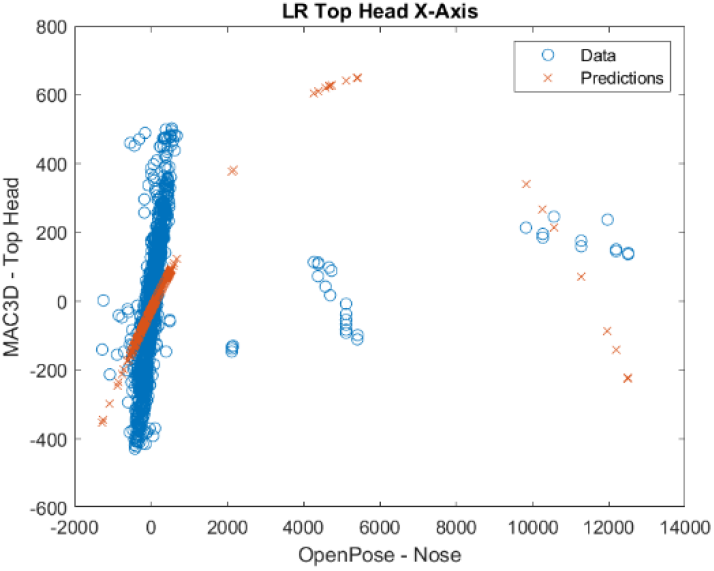

However, as seen in the figure above, a non-linear curving is observed due to the outliers appeared on the right side of the plot. These are OpenPose misdetections. If these erroneous detections are corrected, a relatively straight regression line will also be formed as seen on the left side of the plot.

## IV. Method

Figure 3 shows the proposed framework on how the researchers extracted the 3D points in the world coordinate system. The inputs coming from the four video capture devices were pre-processed and synchronized, as discussed in Section OpenPose was used to extract 2D key points from the pre-processed data while Cortex was used to visualize the MAC3D (.trc) file data. Using the extracted values from the OpenPose 2D data and extracted cortex 3D data, camera calibration was achieved. This has allowed the researchers to recreate the scene and to determine the camera locations and camera parameters even without the conventional checkerboard camera calibration method. With the calibrated multi-camera setup, OpenPose 3D data is available and comparison to the ground truth MoCap data was performed. However, not all key points derived from OpenPose were comparable to the key points present in the MAC3D system. Thus, we objectively selected the key points to be analyzed for comparison. Shown in Figure 4 are the key points that are relatively similar between OpenPose and MAC3D, these points were utilized in the quantitative and qualitative evaluations.

**Fig. 3.**
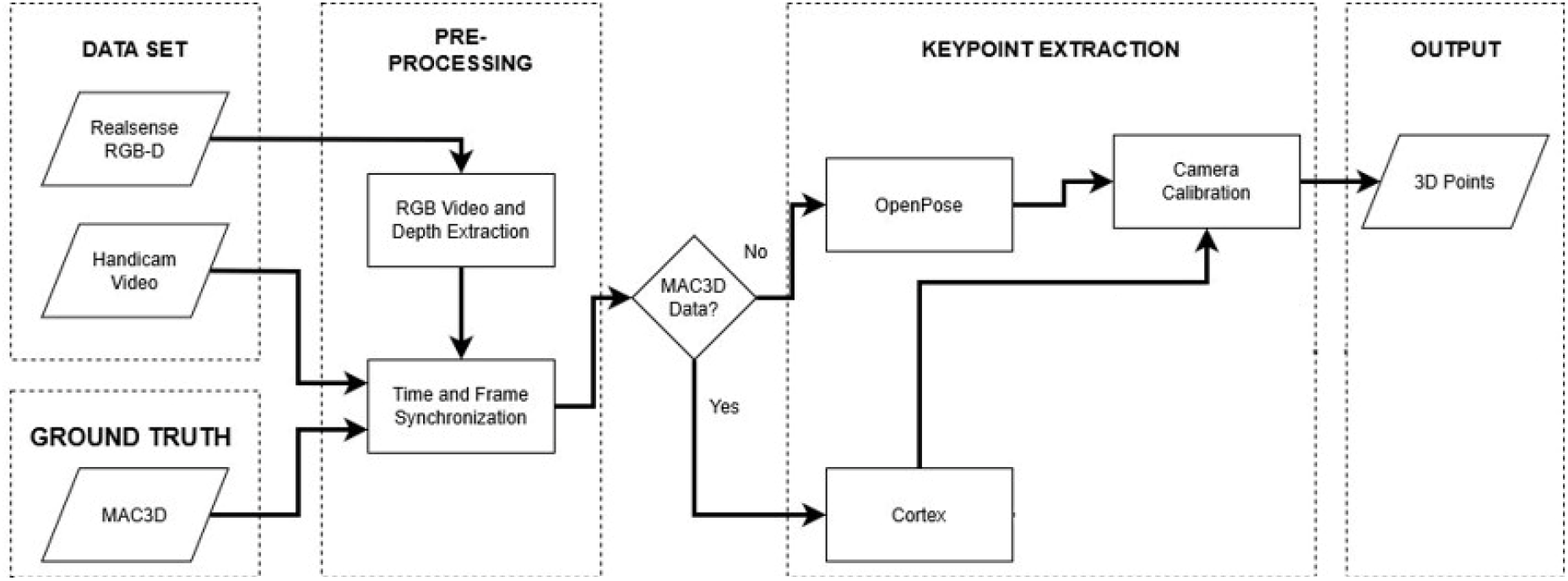
Proposed Pipeline for Evaluating OpenPose 3D with Marker-based Motion Capture System

**Fig. 4.**
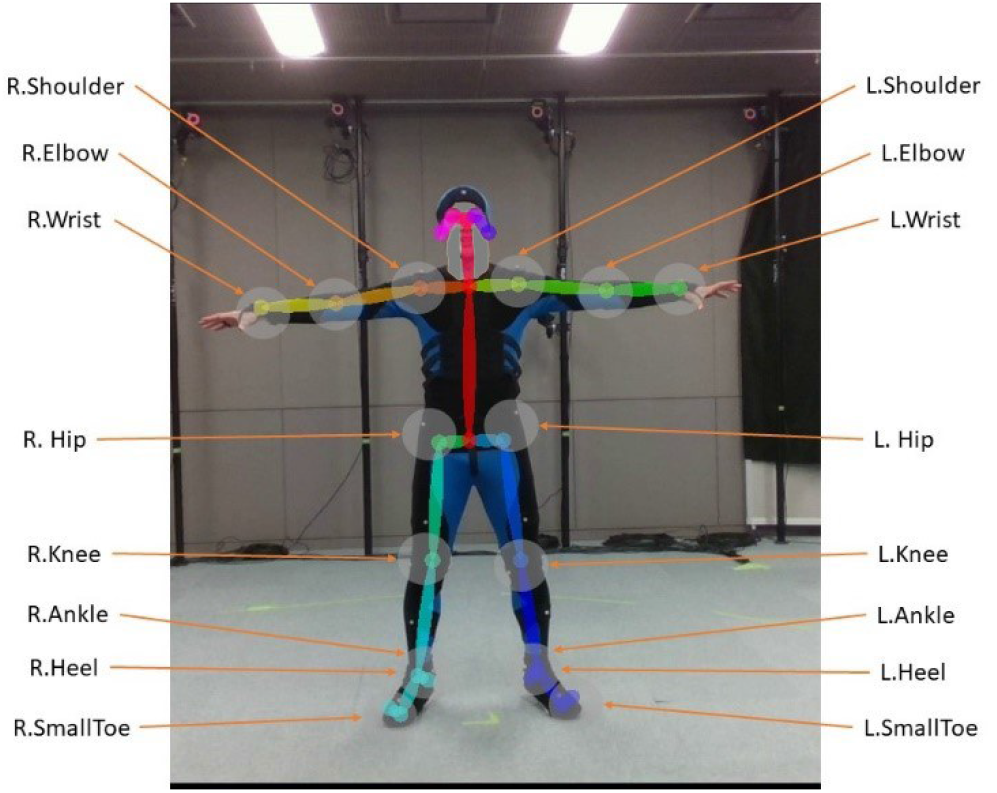
Common points between OpenPose and MAC3D

### A. Overall Framework

The framework has assured the following:

1. RGB videos from three Realsense Camera and the handicam are synchronized.
2. The calibration matrices of each camera are known.
3. Keypoint extraction of OpenPose and MAC3D joint markers have the common base coordinate frame.

### B. OpenPose 3D Triangulation

In order to construct the 3D key points from the 2D key points provided by OpenPose triangulation was utilized. In computer vision, triangulation is the process of determining a point in 3D space given its projection onto two or more images.

In this study, the 2D key points extracted from OpenPose were used to construct the equivalent key points in the 3D space. Making use of the camera calibration, the camera matrix for each recording device was calculated. The function used is the so-called pinhole camera model from OpenCV [9] along with the triangulation method and camera calibration the researchers were able to reproduce the equivalent 3D space key points using the OpenPose 2D key points.

## V. Comparison Results

### A. Quantitative Evaluation

To compare the results predicted by OpenPose with respect to the MoCap data that serves as the ground truth, we used the magnitude of the difference of positions of the key points at the given time Eq. (2): where *v* and *u* are vectors with components *x*, *y*, *z* that represent the coordinates of a keypoint as recorded by the MoCap and predicted by OpenPose respectively.

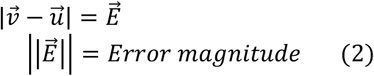

Figure 5 shows the average error of OpenPose of all key points. For the 34.51-sec mark, there is a high error observed due to the dance sequence performed at this time. The dancer moved both his wrists in a criss-cross fashion, making more self-occlusion for the corresponding detections. Figure 6 illustrates the error per keypoint, it is noticeable that the body parts with the highest error are the extremities, especially the wrists.

**Fig. 5.**
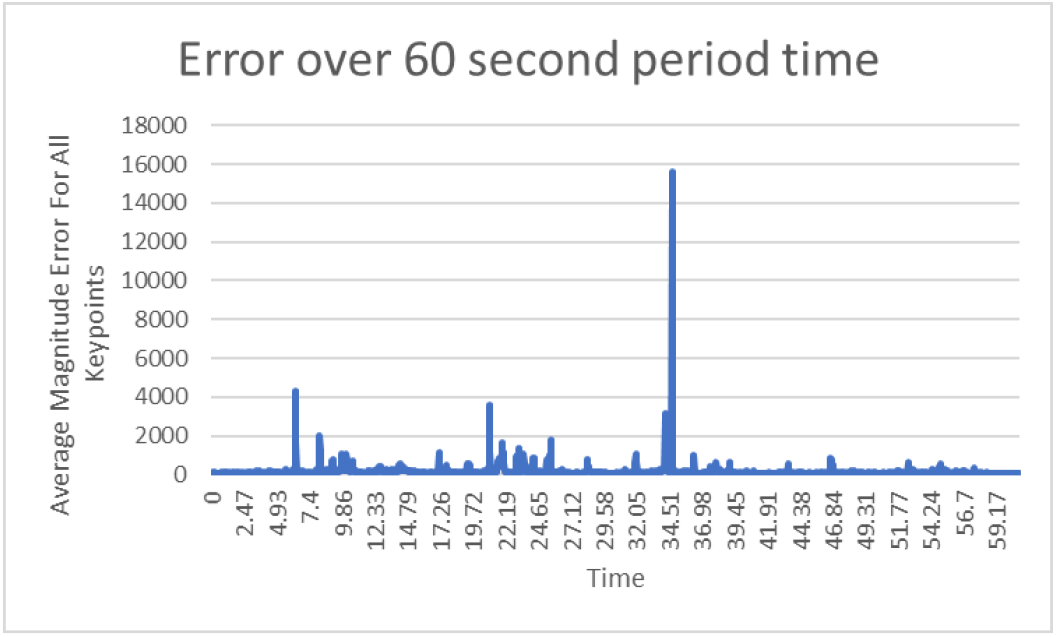
In general, the detection error is less than 2000 mm.

**Fig. 6.**
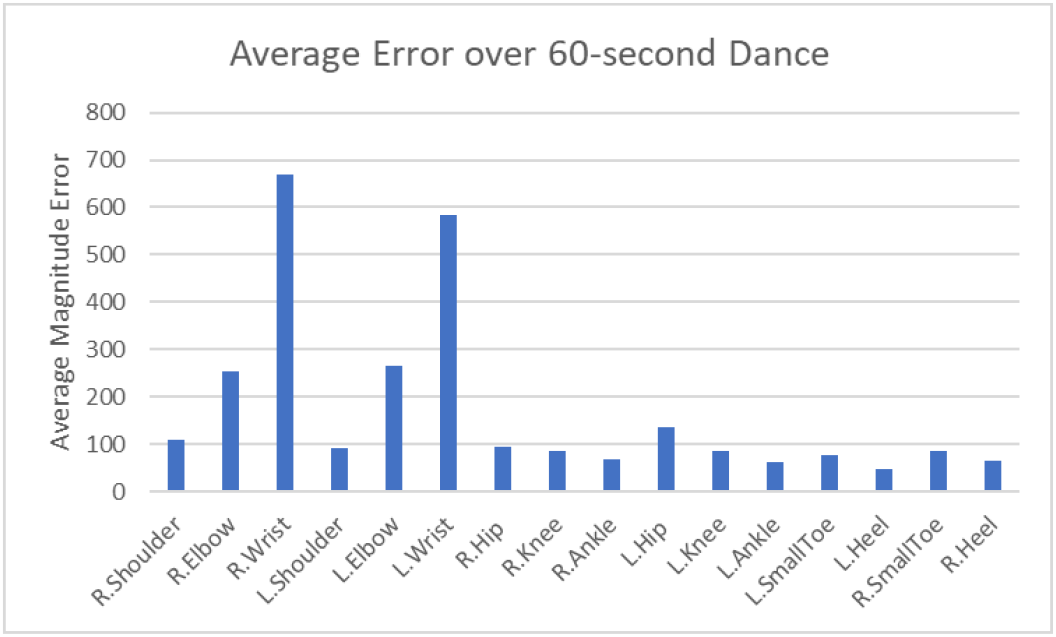
High errors in 3D joint estimated locations occurred on the wrists

**Fig. 7.**
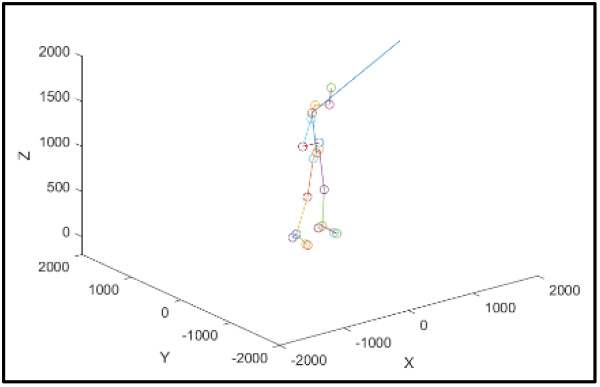
OpenPose misdetection of nose keypoint

### B. Qualitative Evaluation

Samples of success and failed 2D pose snapshots from the OpenPose estimate are illustrated in Figure 8. When the key points are correctly detected on all cameras, the 3D generated pose is expected to be accurate. The markerless pose estimation highly depends on the 2D pose estimates by the OpenPose from each camera. Hence, when the keypoint features extraction was obtained well, the 3D reconstruction is also seen in a proper human skeleton pose. But when there is a mistake on the key points, the 3D skeleton of the dancer is not well reconstructed. Figure 9 depicts the situation for success and failure cases of the OpenPose 3D estimation.

**Fig. 8.**
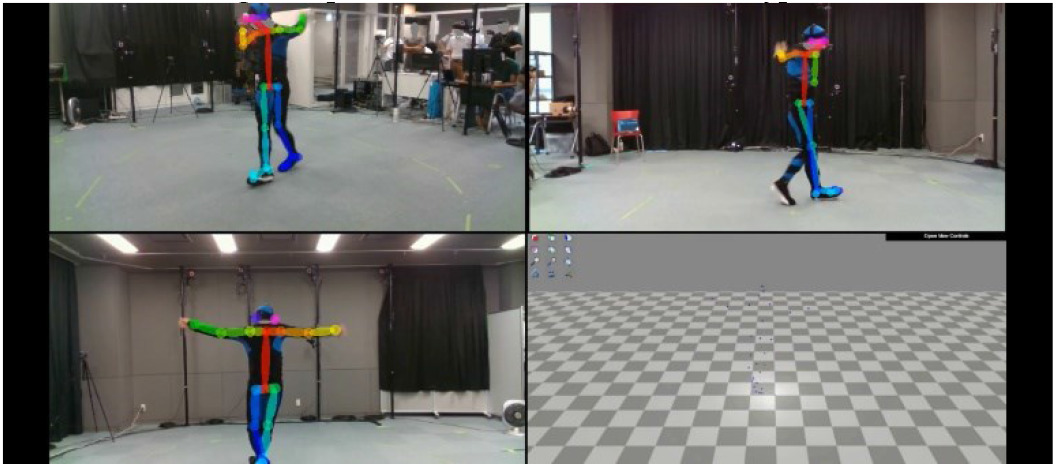
OpenPose 2D Estimates from Three Intel Realsense Camera Positions; Bottom-Right Panel is the visualization of MoCap Data using OpenSim

**Fig. 9.**
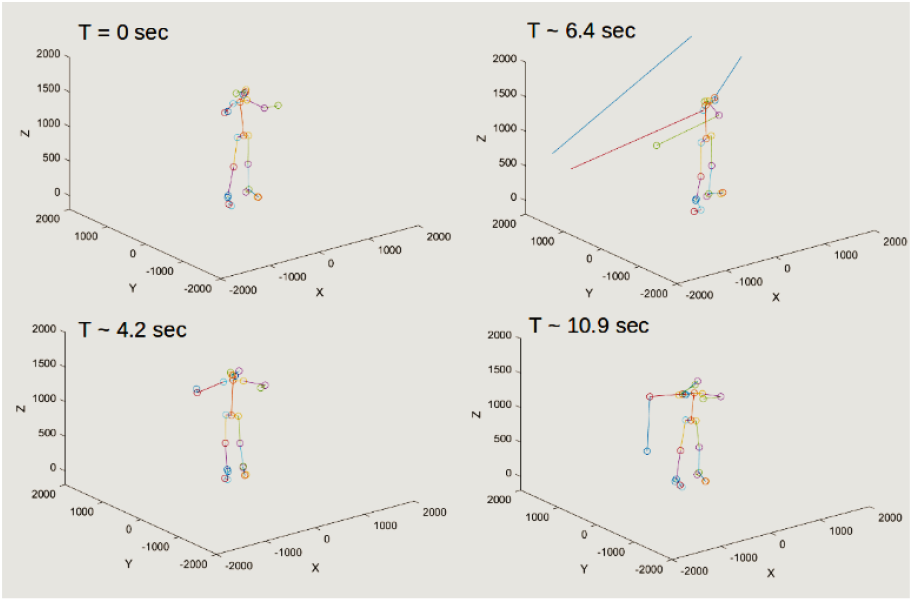
OpenPose 3D Estimates fused from outputs of four videos including handicam; Left panel shows the 3D skeleton in successful scenarios; right panel show where the wrists and elbows were missed due to incorrect 2D.

### C. Discussion on Linear Regression Prediction

In Figure 7, the nose key point was wrongly detected and relatively out of the subject’s body. Since the linear model prediction is based on the OpenPose (Nose), an error in the reconstruction of this particular key point in the world coordinate system would result to an error in the prediction of the MAC3D marker positions (Top, Front, Rear Head).

## VI. Conclusion and Future work

The pop dance motion capture dataset for three Realsense depth sensors, handicam, and MAC3D was synchronized base on the global timestamp. Multi-camera calibration was also done through the use of the marker sets seen at different views of the multiple depth cameras and handicam. Using OpenPose, 3D estimated skeleton pose is generated by fusing the OpenPose estimates on the multiple videos in the synchronized timestamp. However, high upper limb errors on the OpenPose extraction were found when compared to the ground truth motion capture data. To mitigate this, employing a low-pass filter such as Butterworth filter is our future work. Lastly, for the linear model mapping, since the current attempt is an ill-posed problem, in theory, we need to consider two 3D constraints or non-linear mapping. Otherwise, we could also evaluate the case of mapping the relationship between the 16 common selected key points.

## Acknowledgments

The authors would like to thank the Japan Student Services Organization (JASSO) for funding the internship program at Kyushu Institute of Technology. We also acknowledge the participation of Kite Masai, one of the world’s popper (pop dancer) together with Tonyan Kogami, Vinay Kumar, Yousuke Ikeda, and Dr. Shuhei Ikemoto for setting up the motion capture environment. We have a video uploaded in [10] for further information.

